# Haplodiploidy alone does not predict the evolution of eusociality

**DOI:** 10.1101/2025.01.14.633090

**Authors:** Sachin Suresh, Timothy A. Linksvayer

## Abstract

As a corollary to inclusive fitness theory, W.D. Hamilton’s haplodiploid hypothesis proposed that eusociality evolves more readily in haplodiploids because of inflated genetic relatedness between sisters^1^. For the past sixty years, theoreticians have developed models clarifying how haplodiploidy might favor the evolution of eusociality^2–5^. Despite this sustained theoretical interest, the predicted relationship between haplodiploidy and eusociality has only very recently been empirically tested^6,7^. Here, we use two large species-level insect phylogenies with comprehensive sociality and ploidy trait data to test if eusociality evolves more readily in haplodiploids. We find that estimated transition rates to eusociality are higher in haplodiploids, but that this pattern is driven by a much higher transition rate within a single haplodiploid lineage, the aculeate Hymenoptera. Thus, the apparent links between haplodiploidy and eusociality reflect clade-specific dynamics rather than a general effect of haplodiploidy.

## MAIN TEXT

We tested the association between haplodiploidy and eusociality using phylogenetic comparative analyses on the largest available species-level insect phylogenies^8^ and available phenotypic data for sociality and ploidy across insects. Eusociality is traditionally defined by a reproductive division of labor, cooperative brood care, and overlapping generations^5^. Under this definition, eusociality occurs in some aculeate hymenopterans (ants, apid bees, halictid bees, and vespid wasps), termites, thrips, aphids, and a platypodine beetle (Figure 1A). Haplodiploidy (arrhenotoky) is found sporadically across insects, including all hymenopterans and thrips^9^.

**Figure 1.**
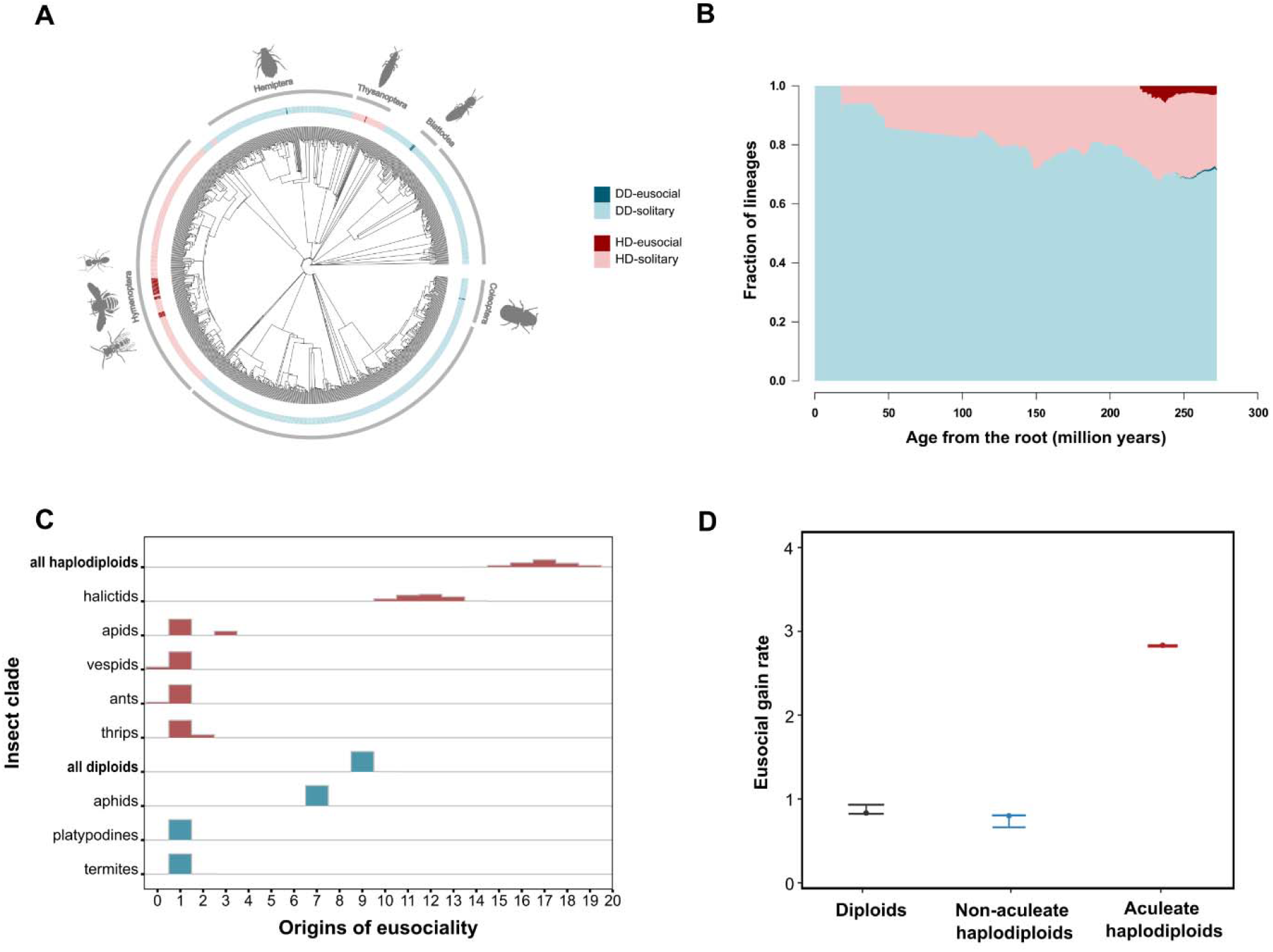
Elevated rates of eusocial evolution are unique to aculeate haplodiploids. A) a representative subset of 1,000 species from the full 68,886 species-level phylogeny of Chesters (2020). The estimated number of independent origins of conventionally defined eusociality is shown in parentheses for each order; B) a lineages-through-time plot. Considering extant species, an estimated 72% of insects are diploid, and 0.96% of these diploids are eusocial. In contrast, 28% of insects are haplodiploid, and 9.66% of these haplodiploids are eusocial. Colors represent four combinations of traits: HD = haplodiploid, DD = diplodiploid (i.e., diploid), EU = eusocial, and non-EU = non-eusocial. C) Distributions of inferred independent origins of eusociality across major insect clades, estimated using stochastic character mapping. Haplodiploid lineages (red) exhibit more eusocial origins than diploid lineages (blue), with most haplodiploid origins concentrated in ants, vespid wasps, and corbiculate bees. (D) Median estimated transition rates from solitary to eusociality under the best-supported model (M_aculeata_), shown separately for diploid lineages, non-aculeate haplodiploid lineages, and aculeate haplodiploid lineages. Error bars indicate 95% confidence intervals across stochastic mappings of ploidy history.

Stochastic character mapping showed that eusociality evolved independently nearly twice as often in haplodiploid lineages (median = 17) as in diploid lineages (median = 9; Figure 1C). Because most insect lineages have been, and remain, diploid (Figure 1B), this concentration of eusocial origins within a smaller portion of the phylogeny implies a higher transition rate from solitary living to eusociality in haplodiploids. Consistent with this, the estimated transition rate to eusociality was approximately seven times higher in haplodiploid than in diploid lineages (Figure S1D).

However, these global comparisons between haplodiploid and diploid lineages do not distinguish whether the observed pattern reflects a general property of haplodiploidy or instead arises from evolutionary dynamics within one or a few clades. Eusociality has evolved repeatedly within the aculeate Hymenoptera (the stinging wasps, bees, and ants) (Figures 1A, 1C, S1C), raising the possibility that this lineage drives the insect-wide association. To test this alternative hypothesis, we fitted regime-dependent models that allowed the transition rate to eusociality to differ among three groups: diploid lineages, haplodiploid lineages outside the aculeate Hymenoptera, and haplodiploid lineages within the aculeate Hymenoptera (Figure 1B, S1B). Under the best-supported model, the transition rate to eusociality was nearly three times higher within aculeate haplodiploid lineages than in both diploid lineages and haplodiploid taxa outside the aculeates (Figure 1B). Model comparison strongly favored this clade-specific model over alternative models lacking an aculeate-specific rate (Figure S1E). We found qualitatively similar results for both phylogenies we used and all combinations of alternative definitions for eusociality and haplodiploidy, including paternal genome elimination that is expected to share many population genetic features with arrhenotoky^9^ (see https://github.com/sachi1n/haplodiploidy-eusociality for complete sets of results).

These results clarify why simple counts of eusocial origins can be misleading. Earlier discussions of the haplodiploid hypothesis have often emphasized the *number* of independent eusocial origins in haplodiploid versus diploid groups^5^. However, since haplodiploid lineages represent only a small fraction of insect diversity (Figure 1B), comparable numbers of origins in haplodiploid and diploid taxa can imply very different transition rates. By estimating transition rates across a species-level phylogeny (Figure S1D) and subsequently considering lineage-specific transition rates (Figure 1B, S1B), our analyses conclude that elevated rates in haplodiploid insects occur within a specific lineage, the aculeate Hymenoptera, rather than across all haplodiploids. This distinction helps reconcile recent empirical results. A family-level comparative analysis reported limited support for a general association between haplodiploidy and eusociality^6^, whereas a recent species-level analysis of 5,678 insects concluded that haplodiploidy has not increased the rate of eusocial evolution^7^. Our study complements and extends this work by using the full taxonomic breadth of a species-level insect phylogeny comprising approximately 69,000 species^8^, with validation on an older tree of nearly 49,000 species. By incorporating all available data on ploidy and sociality traits, our study represents the most comprehensive species-level empirical test of the haplodiploid hypothesis to date, helping to resolve a sixty-year-old debate on the evolutionary association between haplodiploidy and eusociality.

The distribution of eusociality within the Hymenoptera further reinforces the biological interpretation of this conclusion: while all Hymenoptera are haplodiploid, eusociality occurs only within the Aculeata, which represent less than half of the more than 150,000 described hymenopteran species^10^. In hindsight, this pattern may appear straightforward from the distribution of eusociality across insects (Figure 1A). Indeed, despite sustained theoretical attention to the haplodiploid hypothesis and its continued presence in textbooks (Table S1), many social insect researchers have long questioned whether haplodiploidy alone explains eusocial evolution^5^ and have instead emphasized the preponderance of traits specific to aculeate hymenopterans^3^ (Table S1).

Traits such as the sting and specialized nesting behaviors may have favored the evolution of eusociality within the aculeate Hymenoptera^3,5^. However, disentangling the effects of haplodiploidy from traits that are also phylogenetically restricted to specific clades remains challenging. More broadly, eusociality remains rare across insects, including within the aculeate Hymenoptera, and the vast majority of both haplodiploid and diploid species are solitary (Figure 1B). Future studies should seek to clarify the specific traits and ecological, behavioral, and genetic factors that may have facilitated the repeated evolution of eusociality in particular lineages, such as the aculeate Hymenoptera^3^.

## ACKNOWLEDGMENTS

We thank Dr. Liam Revell and Dr. Matt Johnson for their input on the phylogenetic comparative analyses. We thank Dr. Stuart West for the discussion regarding a previous version of our manuscript. We also thank Juergen Liebig and other members of the Social Insect Research Group at ASU for feedback on the study and members of the Linksvayer lab for feedback on the manuscript. This work received partial support from NSF IOS 2128304 to TAL.

## AUTHOR CONTRIBUTIONS

SS and TAL conceived and designed the research; SS collected the data; SS and TAL analyzed the data; SS and TAL wrote the paper.

