## Supplemental file for "Haplodiploidy alone does not predict the evolution of eusociality"

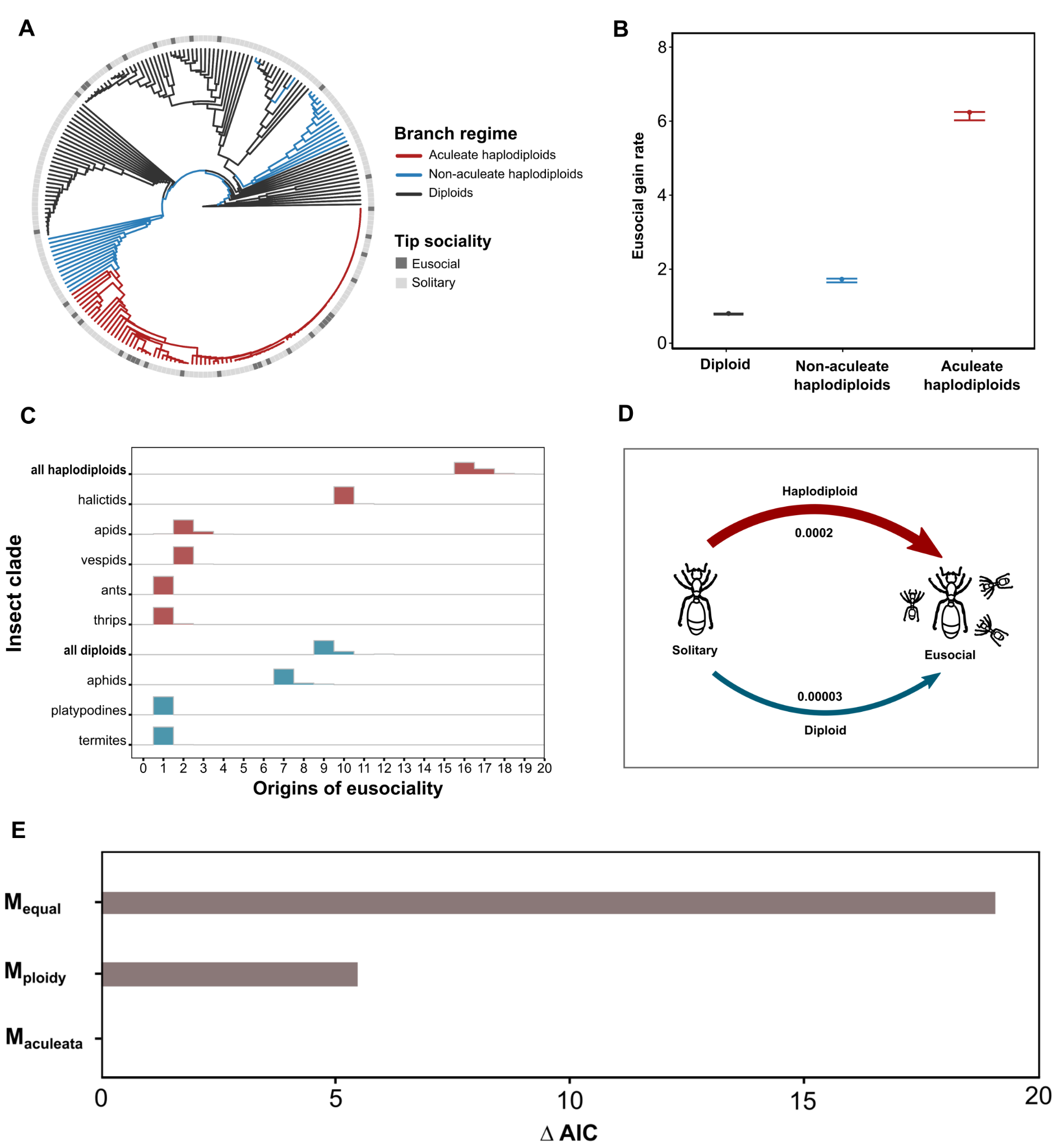

**Figure S1. Elevated rates of eusocial evolution in aculeate haplodiploids.** (A) Collapsed Chesters 2017 phylogeny used for the regime-dependent comparative analyses. To reduce redundancy from large clades lacking variation in ploidy or sociality, monomorphic clades for both traits were pruned to a single representative lineage before analysis. Branch colors indicate inferred ploidy regime based on stochastic character mapping: diploid lineages (black), non-aculeate haplodiploid lineages (blue), and aculeate haplodiploid lineages (red). Tip shading indicates the extant sociality state (eusocial vs. solitary). (B) Median estimated transition rates from solitary to eusocial states under the best-supported model (M_aculeata_), shown separately for diploid lineages, non-aculeate haplodiploid lineages, and aculeate haplodiploid lineages for Chesters 2017 phylogeny. Error bars indicate 95% confidence intervals across stochastic mappings of ploidy history. (C) Distributions of inferred independent origins of eusociality across major insect clades for Chesters 2017 phylogeny, estimated using stochastic character mapping. Haplodiploid lineages (red) exhibit more eusocial origins than diploid lineages (blue), with most haplodiploid origins concentrated in ants, vespid wasps, and corbiculate bees. D) Estimated transition rates from solitary to eusociality across the whole tree in haplodiploid lineages relative to diploids for Chesters 2020 phylogeny. The conventional definition of eusociality and the strict definition of haplodiploidy are used to estimate origins and transition rates. (E) Model comparison based on median ΔAIC across stochastic mappings, where ΔAIC represents the difference in Akaike Information Criterion between each model and the best-supported model (ΔAIC = 0). Lower ΔAIC values indicate stronger support for a model. A model allowing distinct eusociality gain rates in aculeate haplodiploids, non-aculeate haplodiploids, and diploids (M_aculeata_) is strongly favored over models assuming equal rates across lineages (M_equal_) or rates differing only by ploidy (M_ploidy_).

| **Excerpts from biology textbooks presenting the haplodiploidy hypothesis** | |
| --- | --- |
| **Book** | **Quote(s) on the haplodiploid hypothesis** |
| Alcock, J. (2013). Animal Behavior: An Evolutionary Approach, 10th ed. (Sinauer Associates). | Hamilton realized that the coefficient of relatedness  between full sisters in hymenopteran species could be higher than the 0.5 figure that applies to full siblings in most other organisms. A higher coefficient of relatedness among sisters could make adaptive altruism among sisters more likely to evolve, thereby explaining why sterile female workers were especially well represented among the Hymenoptera |
| Nordell, S.E., and Valone, T.J. (2017). Animal Behavior: Concepts, Methods, and Applications, 2nd ed. (Oxford University Press). | Hamilton suggested that haplodiploidy favors the evolution of sterile daughters because they can achieve higher fitness by helping to raise sisters instead of their own daughters |
| Krebs, J.R., Davies, N.B., and Parr, J. (1993). An Introduction to Behavioural Ecology, 3rd ed. (Blackwell Scientific Publications). | W.D. Hamilton (1964) was the first to fully appreciate the significance of the special genetic predisposition of Hymenoptera to form sterile castes.  In other words, because of haplodiploidy full sisters are more closely related to one another than are parents and offspring in a normal diploid species. Hymenopteran queens are diploid and are therefore related to their sons and daughters by the usual 0.5. A sterile female worker can therefore make a greater genetic profit by rearing a reproductive sister than she could if she suddenly became fertile and produced a daughter! This extraordinary state of affairs also suggests why in the Hymenoptera only females help to rear sisters |
| Bergstrom, C.T., and Dugatkin, L.A. (2016). Evolution, 2nd ed. (W. W. Norton & Company). | Why is Hymenoptera particularly prone to evolve eusociality?  A genetic relatedness of 0.75 between sisters has the remarkable effect of making females more related to their sisters than to their own offspring. The answer may lie, in part, in the unusual genetic architecture of the hymenopterans.  Because of the asymmetries in genetic relatedness, we predict that eusocial behaviors should be displayed by female workers but not by males (drones). |
| Freeman, S., and Herron, J.C. (2007). Evolutionary Analysis, 4th ed. (Pearson Prentice Hall). | William Hamilton (1972) proposed that these taxa are predisposed to eusociality by their unusual form of sex determination.  Hamilton argued that females can maximize their inclusive fitness by acting as workers and investing in the production of sisters, rather than by acting as reproductives. |
| **Theoretical studies linking ploidy and the evolution of sociality** | |
| **Study** | **Major finding(s)** |
| Trivers and Hare (1976) | The asymmetrical relatedness in haplodiploid species predisposes daughters to evolve eusocial behavior, provided they can exploit these asymmetries by either producing more females than the queen would prefer or by gaining some or complete control over male production genetics. |
| Charlesworth (1978) | In haplodiploids, both kin selection and parental manipulation can establish an evolutionarily stable strategy, resulting in an intermediate frequency of altruists within a sibship. When the cost/benefit ratio exceeds a critical value, selection typically pushes the fitness sacrificed by the altruists to its maximum, thereby increasing the fitness of their siblings. This mechanism may help to explain the origin of eusociality |
| Charnov (1978) | The genetic asymmetry in haplodiploids predisposes daughters to evolve eusocial behavior, provided they can bias the sex ratio or selectively substitute sons for brothers |
| Craig (1979) | Haplodiploidy promotes the evolution of eusociality through parental manipulation. Since male offspring cost less to produce (haploid), manipulating daughters into workers benefits the queen even when brood efficiency is slightly reduced. If the cost of forcing offspring to be sterile workers reduces their brood-raising efficiency by less than half, the manipulation allele will still spread because the queen’s genetic gain outweighs the loss |
| Aoki and Moody (1981) | Haplodiploidy enables workers to enhance their inclusive fitness by substituting their own sons for brothers. This ability makes haplodiploidy more conducive for the evolution of eusocial behavior, particularly under conditions where worker traits and discriminatory behavior are tightly linked genetically. |
| Iwasa (1981) | Eusociality evolves more easily in haplodiploids because the threshold benefit-to-cost ratio for altruistic behavior is initially lower in haplodiploid systems than in diploids. |
| Seger (1983) | Partial bivoltinism can create conditions that favor the evolution of eusociality through alternating sex-ratio biases in haplodiploids |
| Grafen (1986) | Haplodiploidy facilitates eusociality primarily through asymmetry in genetic relatedness and the ability to produce and manipulate split sex ratios. These mechanisms make sib rearing more advantageous in haplodiploids |
| Godfray and Grafen (1988) | In haplodiploids, ability of unmated females to produce viable male offspring leads to a natural variation in sex ratios within populations. This variation creates conditions where sib rearing is advantageous, particularly in female-biased broods of mated females, thereby facilitating the evolution of eusociality without the need for initial sex discrimination by workers. |
| Ratnieks (1988) | Haplodiploidy facilitates eusociality via cooperation enforcement with worker policing |
| Reeve (1993) | The evolution of eusociality is favored in haplodiploids due to a protected invasion effect: Dominant alleles for maternal and female alloparental care are more resistant to loss compared to paternal-care alleles in haplodiploids. |
| Wade (2001) | Haplodiploidy facilitates eusociality through the evolution of maternal care and maternal effect genes that produce antagonistic pleiotropic effects on sons, leading to female-biased sex ratios and enhanced kin selection dynamics. |
| Linksvayer and Wade (2005) | Eusociality may more readily evolve in haplodiploids because eusociality is derived from subsociality, and subsociality may more readily evolve in haplodiploids. |
| Boomsma (2007) | Haplodiploidy and strict monogamy facilitates the evolution of eusociality |
| Fromhage and Kokko (2011) | Haplodiploidy and monogamy synergistically facilitate the evolution of eusociality by enhancing relatedness among siblings and promoting positive phenotypic assortment. This synergy allows eusocial colonies to achieve greater efficiency and resilience, thus supporting the evolution and maintenance of complex social structures. |
| Johnstone et al. (2011 | Haplodiploidy facilitates eusociality by increasing the local relatedness among females through male-biased dispersal. This higher relatedness enhances the kin-selected benefits of helping behaviors, making it more likely for altruistic alleles to spread in haplodiploid populations. |
| Gardner et al. (2012) | Haplodiploidy can promote eusociality under certain conditions, primarily through split sex ratios caused by queen virginity or replacement; the overall effect is likely small. Monogamy appears to have been a more critical factor in the evolution of eusociality. |
| Gardner and Ross (2013) | Haplodiploidy facilitates eusociality primarily by enabling precise sex-ratio adjustments, which favor the evolution of helping behaviors in sex-biased systems. This mechanism operates through local resource enhancement and the rarer-sex effect, creating a feedback loop that promotes altruism and cooperative behaviors. |
| Alpedrinha et al (2013) | Haplodiploidy facilitates eusociality primarily through the effects of split sex ratios and worker reproduction. |
| Rautiala et al. (2014) | Haplodiploidy facilitates eusociality primarily through relatedness asymmetries and the ecological dynamics of unmated females leading to split sex ratios. |
| Davies et al. (2016) | Relatedness asymmetry in haplodiploids plays a role in facilitating the evolution of eusociality. However, the ecology of sex, including preadaptation for certain traits, sex-ratio adjustment, and the specifics of the mating system (e.g., monogamy and sib-mating), significantly shapes the patterns of helping in arthropod societies. |
| Quinones and Pen (2017) | Haplodiploidy facilitates eusociality through genetic relatedness but also significantly through maternal sex ratio adjustment |
| Rautiala et al. (2019) | Haplodiploidy promotes the evolution of female helpers by lowering the benefit threshold for helping, regardless of the population sex ratio. |
| Kennedy and Radford (2020) | Genetic predisposition of supersister relatedness in haplodiploids combined with the greater impact of sibling quality on female fitness can drive the evolution of altruistic behaviors. |

**Table S1. Representative excerpts from biology textbooks and summaries of theoretical studies linking ploidy and sociality**. Summary of the representative textbook descriptions of the haplodiploidy hypothesis and an overview of recent theoretical studies examining the relationship between ploidy and the evolution of sociality. This table builds in part on the summary presented in Table 1 of Joshi and Wiens (2023), with additional sources and quotations included to reflect the range of theoretical work on the haplodiploidy hypothesis.

**Supplemental Experimental Procedures**

**Insect phylogeny**

We used two species-level insect phylogenies: Chesters 2017^S^^1^, containing 49,358 species, and Chesters 2020^S^^2^, with 68,886 species. To our knowledge, the 2020 tree is the most comprehensive species-level insect phylogeny currently available. We time-scaled these trees using treePL, which implements a penalized-likelihood algorithm to estimate divergence time^S^^3^. We included minimum age constraints from twenty-two fossils^S^^4^.

**Trait data**

We collected ploidy data by conducting a literature review and using the online Tree of Sex database^S^^5^. In our trait data, there were two forms of haplodiploidy: 1) a “stricter” definition of haplodiploidy that only considers arrhenotoky, which includes the large majority of cases and has been the conventional focus of discussions of haplodiploid insects. Sociality data was manually curated from existing literature^S^^6–S15^. 2) a “broader” definition of haplodiploidy, which includes both arrhenotoky and paternal genome elimination^S^^16^, Our trait data also includes two definitions of eusociality: 1) a “conventional” eusocial definition, characterized by the reproductive division of labor, cooperative care of the brood, and overlapping generations^S^^17,S18^; 2) a “superorganismal” definition of Boomsma and Gawne^S^^19^ characterized by irreversible worker caste differentiation. Following their classification, we consider ants (Formicidae), higher termites (Termitidae), corbiculate bees (Apini, Meliponini, Bombini), and vespine wasps (Vespinae) as superorganismal. Eusociality, by the conventional definition, occurs in five insect orders (Thysanoptera, Hemiptera, Blattodea, Hymenoptera, and Coleoptera), and the strict definition of haplodiploidy (arrhenotoky) only co-occurs with eusociality in the insect orders Thysanoptera and Hymenoptera (Fig.1a).

**Stochastic Character Mapping of Eusocial Origins**

To estimate the number of independent evolutionary origins of eusociality under different ploidy regimes, we used stochastic character mapping (SCM), a simulation-based method for reconstructing discrete trait histories on a phylogeny^S^^20^, as implemented in the R package phytools^S^^21^. This approach generates a posterior distribution of character histories by simulating transitions along each branch, conditioned on observed tip states and a specified model of evolution. We modeled eusociality as a binary trait (0 = non-eusocial, 1 = eusocial), applying the all-rates-different (ARD) model of trait evolution, which allows the rate of transition from 0 to 1 to differ from the rate of transition from 1 to 0. This model is the most general two-state Markov model and avoids making restrictive or potentially unjustified assumptions about reversibility^S^^22^. The ARD framework is particularly suitable for traits like eusociality, where the evolutionary gain and loss may occur under different ecological or genetic constraints. To specify priors on the ancestral state of eusociality, we used the “fitzjohn” approach^S^^23^, which derives the root state prior from the stationary distribution of the fitted transition matrix. This method is advantageous in avoiding arbitrary assumptions about trait polarity, especially when the true ancestral state is uncertain or when multiple gains and losses are expected. We conducted 1,000 replicate stochastic maps for each pairwise combination of trait definitions (strict vs. broad eusociality and strict vs. broad haplodiploidy). For each replicate, we recorded the number of independent origins of eusociality and categorized these transitions by whether they occurred on diploid or haplodiploid branches. This approach enabled us to generate robust estimates of the frequency of eusociality origins associated with each ploidy regime while incorporating uncertainty in both the trait reconstruction and the phylogenetic relationships.

**Testing for Correlated Evolution**

To formally assess whether the evolution of eusociality is statistically associated with ploidy type, we used Pagel’s test of correlated evolution for binary traits^S^^22^, as implemented in the *fitPagel()* function in *phytools*. This method models the joint evolution of two traits under a continuous-time Markov process and compares alternative models reflecting different assumptions about their dependence. We fit four models for each trait combination: (1) a model in which eusociality and haplodiploidy evolve independently (4 rate parameters), (2) a model in which the evolution of haplodiploidy depends on social state (6 parameters), (3) a model in which the evolution of eusociality depends on ploidy (6 parameters), and (4) a fully dependent model in which transitions in each trait depend on the state of the other (8 parameters). All models were run using the ARD rate structure, allowing asymmetric transition rates. As with the stochastic mapping analysis, we used the “fitzjohn” prior on root states. Model fit was evaluated using Akaike Information Criterion (AIC), and models were compared using ΔAIC thresholds, where ΔAIC > 2 was taken as evidence for better support of the more complex model^S^^24^. For the best-fitting model, we extracted the estimated transition rates to quantify the relative likelihood of eusociality evolving under haplodiploid versus diploid regimes.

**Regime-dependent models of eusocial evolution**

To examine heterogeneity in eusociality transition dynamics among haplodiploid taxa, we conducted an additional comparative analysis in which the rate of change from solitary to eusocial states was estimated separately for diploid lineages, haplodiploid lineages outside Aculeata, and haplodiploid lineages within Aculeata. The phylogeny was first filtered to include only taxa with complete ploidy and sociality information. To minimize redundancy arising from large clades lacking variation in either trait, clades that were invariant for both ploidy and sociality were reduced to a single representative lineage before analysis. Uncertainty in the evolutionary history of ploidy was incorporated using stochastic character mapping^S^^20^. Haplodiploidy (diploid vs. haplodiploid) was mapped onto the collapsed phylogeny using an equal-rates model, generating multiple stochastic realizations of ploidy histories. These mappings partitioned each branch into segments assigned to diploid or haplodiploid regimes. Aculeate Hymenoptera were defined as a monophyletic clade based on the most recent common ancestor of extant aculeate families, and branch segments were classified according to whether they fell within or outside this clade. For each mapped ploidy history, we fit continuous-time Markov chain models^22^ describing transitions between solitary and eusocial states. The model was parameterized by an origin rate (q01) and a loss rate (q10), with q01 allowed to vary among regimes and q10 shared across regimes. Transition probabilities along each branch were computed using matrix exponentiation, and likelihoods were calculated using a pruning algorithm for discrete traits. Model parameters were estimated by maximum likelihood with multiple random starting values to avoid local optima. We compared three nested models: a model with no regime dependence, a model in which transition rates differed between diploid and haplodiploid lineages, and a model in which haplodiploid lineages inside and outside Aculeata were assigned distinct transition rates. Model support was assessed using Akaike Information Criterion (AIC), and likelihood-ratio tests were used to compare nested models. Parameter estimates and model support were summarized across stochastic maps to account for uncertainty in ploidy history. All statistical analyses were performed using the *phytools* package^S^^25^ in R 4.3.2^S^^26^. All data and analysis codes used for this study are available on GitHub (<https://github.com/sachi1n/haplodiploidy-eusociality>)
